# Development of an EMT-related exosomal miRNA signature that can predict prognosis in hepatocellular carcinoma

**DOI:** 10.1101/2025.09.06.674651

**Authors:** Olka Missaghi, Leman Nur Nehri, Sepehr Bakhshi, Oğuzhan Karaosmanoğlu, Hülya Sivas, Aybar Can Acar, Sreeparna Banerjee

## Abstract

Chemoresistance and epithelial-mesenchymal transition (EMT) are associated with failure of cancer chemotherapy and poor survival of patients. We have previously shown that chemoresistance and stemness in hepatocellular carcinoma (HCC) cells was accompanied by the development of partial EMT (p-EMT) and identified a number of EMT-associated proteins that are released from exosomes. In this study, we aimed to identify and classify the differentially expressed (DE) exosomal miRNAs from chemoresistant HuH7 cells undergoing p-EMT. Out of the fifty-four miRNAs that were enriched in the exosomes from these cells compared to controls, thirteen were identified in the exosomes isolated from the serum of HCC patients. These miRNAs targeted genes that were associated with cell-cell junctions, extracellular matrix, cytoskeleton, transcription and signal transduction. Univariate Cox regression analysis indicated that 11/13 miRNAs were associated with either favorable (n=4) or worse (n=7) prognosis. A machine learning algorithm indicated that seven miRNAs (miR 215-5p, miR 340-5p, miR 210-3p, miR 19a-3p, miR19b-3p, miR 1266-5p and miR 25-3p) could predict worse prognosis in multiple datasets with 64-68% accuracy. A Bayesian Inference network analysis with the thirteen miRNAs and their key target proteins, along with EMT and survival as the nodes indicated that the common denominator was transcription, suggesting that the exosomal miRNAs released from cells undergoing p-EMT can mediate phenotypic changes in cells through transcriptional regulation.

## 1. Introduction

Epithelial to mesenchymal transition (EMT) is a process during which epithelial cells lose their basal-apical polarity, display weakened cell-cell junctions and acquire migratory and invasive (mesenchymal) traits. EMT is known to increase motility, stemness characteristics and drug resistance in cancer (Najafi et al., 2020; O’Reilly et al., 2019). However, there is considerable heterogeneity in the gain of mesenchymal traits suggesting that EMT is not a binary state, with most cells displaying a hybrid epithelial/mesenchymal phenotype. In hybrid/partial EMT (p-EMT), cells can express both epithelial and mesenchymal markers such as E-cadherin and Vimentin and may migrate or circulate as tumor cell clusters (Subbalakshmi et al., 2022). These cells may also acquire stem cell features such as drug-resistance, apoptosis-resistance, and enhanced self-renewal in addition to migration and invasion (Pan et al., 2021) .

p-EMT is a closely regulated process. Transcription factors such as SLUG and ΔNP63α were shown to be associated with a p-EMT phenotype in basal like breast cancer (Subbalakshmi et al., 2022). However, recent data suggest that post-transcriptional mechanisms such as alterations in subcellular localization of epithelial proteins (Aiello et al., 2018) or activation of non-coding RNAs such as microRNAs can be involved in the process of p-EMT (Saitoh, 2018). MicroRNAs primarily regulate protein expression at the post-transcriptional level by binding to the 3’-untranslated regions (UTR), 5’UTR, or to the coding region of mRNAs and have been reported to modulate EMT in cancer (Lou et al., 2018).

Exosomes are small vesicles that originate inside multivesicular bodies. These vesicles have an average diameter of 50-150 nm and can carry functional molecules such as non-coding RNAs including miRNAs, lncRNA, DNA, mRNA and a large repertoire of proteins (Baig et al., 2020). Primary tumor cells releasing exosomes may communicate with cells in the immediate tumor microenvironment or at distant sites via the release of encapsulated RNA molecules (coding or non-coding) that may enhance cancer progression (Lucotti et al., 2022). For example, incubation of hepatocellular carcinoma (HCC) cells with exosomes containing miR-92a-3p was shown to promote EMT by targeting the tumor suppressor PTEN and activating downstream signaling via the PI3K/AKT pathway (Yang et al., 2020). RNA binding proteins can mediate the encapsulation of non-coding RNAs into exosomes via the autophagy related protein LC3 (Leidal et al., 2020). Additionally, the release of exosomes was shown to be enhanced by EMT, suggesting a role in the progression of cancer (Tauro et al., 2013). Some tumor-derived exosomal microRNAs may also inhibit EMT. For example, miR-375-3p was shown to target the mesenchymal transcription factor YAP1 in prostate cancer and promote mesenchymal to epithelial transition (MET) (Selth et al., 2017). The loading of miRNAs to exosomes can also be highly regulated and selective, emphasizing their diagnostic and prognostic value in cancer due to their ease of collection and high stability.

We have previously shown that HCC cells undergoing p-EMT released exosomes enriched in EMT related protein such as fibronectin 1 (FN1), collagen type II alpha 1 (COL2A1) and native fibrinogen gamma chain (FGG) (Karaosmanoğlu et al., 2018). In the current study, the small RNA content of the exosomes released from the same HCC cells undergoing p-EMT or control cells was isolated, sequenced and validated. Bioinformatics analyses indicated that the exosomal miRNAs from HCC cells undergoing p-EMT were of prognostic significance across multiple datasets and could regulate important cellular processes such as cell-cell adhesion, cytoskeletal network, signal transduction and transcription. A Bayesian Inference network analysis using EMT and survival as nodes indicated that the common denominator for the 13 miRNAs and the primary cellular function that they regulated was transcription. Thus, exosomal miRNAs released from cancer cells undergoing p-EMT have a high probability of mediating phenotypic changes via the regulation of transcription.

## 2. Materials and methods

### 2.1 Cell lines and culture conditions

We have previously reported the stable overexpression of Slug in the HCC cell line Huh7 (Karaosmanoğlu et al., 2018). These cells showed characteristics of partial EMT via the upregulation of both E-cadherin and Vimentin and chemo-resistance compared to the empty vector transfected control cells (EV). Both Slug overexpressing and control cells were cultured in DMEM (Sigma-Aldrich, St. Louis, MO, USA) supplemented with 250 μg/mL G418 (Sigma-Aldrich, St. Louis, MO, USA) for the maintenance of overexpression, 10% fetal bovine serum (Sigma-Aldrich, St. Louis, MO, USA), 100 U/mL penicillin and 100 μg/mL streptomycin (Sigma-Aldrich, St. Louis, MO, USA) at 37 °C in a humidified atmosphere containing 5% CO_2_.

### 2.2 Exosome and exosomal small RNA isolation

Slug overexpressing and control (EV) cells were seeded as 2.5x10^5^/ 25 cm^2^ flask, and serum-starved after reaching 70-80% confluency. Conditioned medium was collected after 24 hours of incubation and exosomes were isolated according to the manufacturer’s instructions (Total Exosome Isolation kit, Invitrogen, California, USA). Briefly, the conditioned medium was centrifuged at 2000 × *g* for 30 minutes to remove cells and debris. Next, two volumes of the supernatant were mixed with one volume of Total Exosome Isolation reagent and incubated overnight at 4 °C. Next, the samples were centrifuged at 10,000 × *g* for 1 hour at 4°C. The pellet, which contained the exosomes, was dissolved in a lysis buffer from the Nucleospin miRNA kit (Macherey Nagel). Small RNAs were isolated according to the protocol of the manufacturer.

### 2.3 Small-RNA sequencing and data analysis

Small RNA isolated from Slug overexpressing and control cells were sequenced at Fasteris SA, Plan-les-Ouates, Switzerland. Briefly, libraries were prepared with NEB Next Small RNA Library Prep Set for Illumina (NEB, Ipswich, UK) according to the manufacturer’s protocol. Then, single-read sequencing with 1 × 50 bp read length using the Hiseq 2000 platform (Illumina, San Diego, CA, USA) was applied. miRNA read counts per library were normalized for the comparison of the expression of exosomal miRNAs released from Slug overexpressing and control cells. Next, the miRNA read counts were analyzed for differences using the R-package DESeq (Love et al., 2014).

### 2.4 Validation of miRNA expression by quantitative RT-PCR

Exosomal small RNAs were isolated from Slug overexpressing and control cells as described above. Among the significantly upregulated miRNAs from the small RNA sequencing, three miRNAs (miR-101-3p, miR-296-3p and miR-885-5p) were selected randomly and their levels were determined by quantitative RT-PCR using a specific and quantitative method as described previously (Balcells et al., 2011). The expressions of miR-16-1-3p, miR-1228-3p were used as internal control as these miRNAs are known to have stable expression in HCC (Hu et al., 2014; Kelly et al., 2015). For the generation of cDNA from the RNA, 10 ng of small RNA, 1 μl of 10X poly(A) polymerase buffer, 0.1 mM of ATP, 1 μM of RT-primer, 0.1 mM of each deoxynucleotide, 100 U of MuLV reverse transcriptase (New England Biolabs, USA) and 1 U of poly(A) polymerase (New England Biolabs, USA) were mixed in a PCR tube. Next, the small RNAs were converted to cDNA with incubation at 42°C for 1 hour followed by inactivation of the enzyme at 95°C for 5 minutes. Quantitative RT-PCR was carried out with 1 μl of 10X diluted cDNA, 250 nM of each primer (Table 1), 5 μl of Brilliant SYBR Green qPCR master mix (Agilent Technologies, Palo Alto, CA, USA), in a real-time PCR platform (Mx3005P, Agilent Technologies, Palo Alto, CA, USA).

**Table 1.**
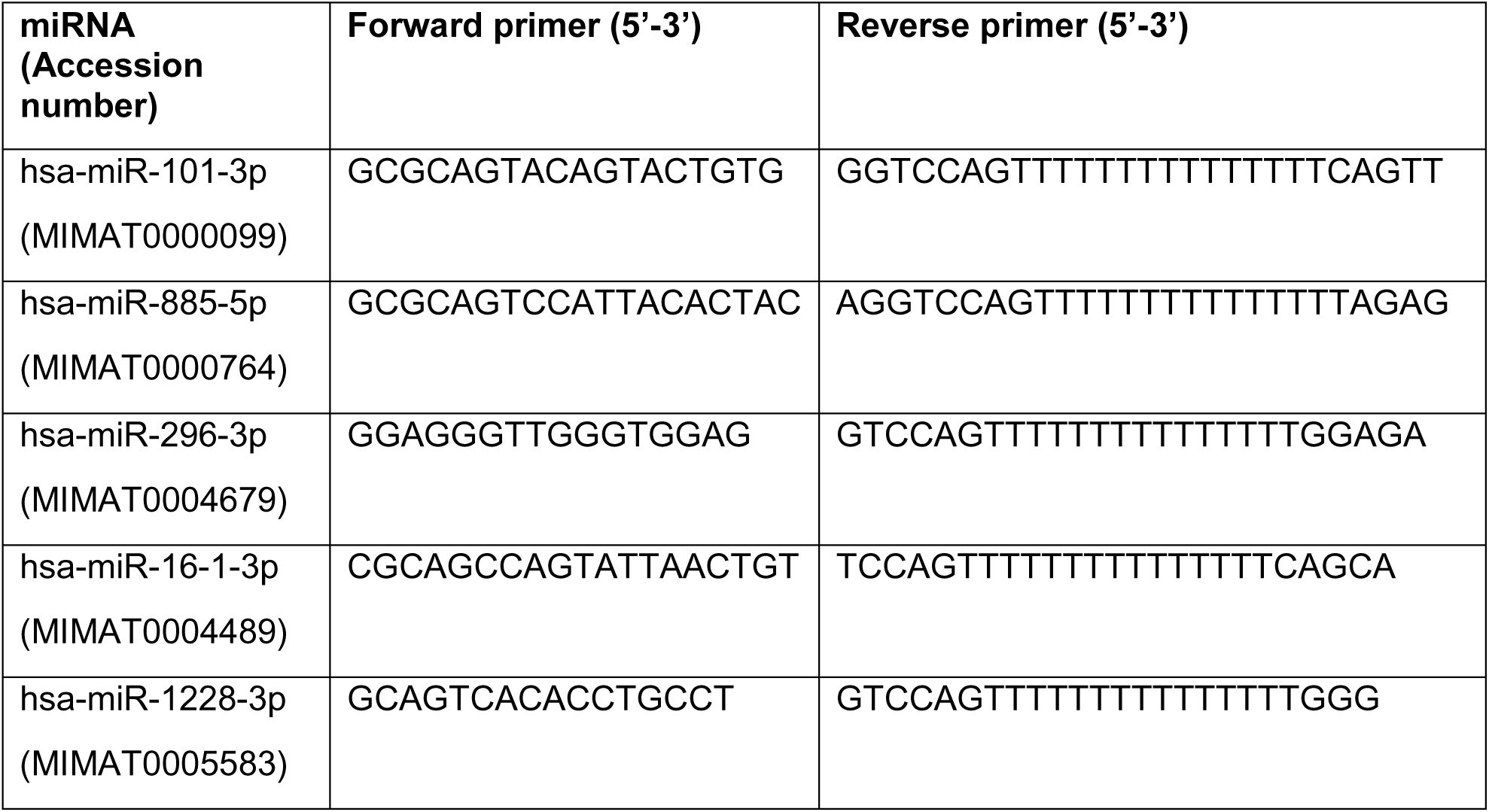
The primer sequences used for miRNA qRT-PCR.

### 2.5 Bioinformatics analyses

#### 2.5.1 Data Validation

To determine whether the miRNAs identified with the small RNA sequencing from the Slug overexpressing cells were relevant to human HCC, we examined miRNA expression datasets of HCC tissues with survival data from the National Center for Biotechnology Information (NCBI) online Gene Expression Omnibus (GEO) database (https://www.ncbi.nlm.nih.gov/geo). The search strategy used was "carcinoma, hepatocellular" OR hepatocellular carcinoma AND "micrornas" OR miRNA AND exosome AND “neoplasm metastasis" OR metastasis. Only 2 datasets were found in the initial search: GSE209611 (Kawamura et al., 2022) and GSE106452 (Fang et al., 2018). Our inclusion criterion was the availability of sequencing data from exosome derived miRNAs from HCC tumor and benign samples. After carefully screening the available samples, we selected 14 samples from GSE209611 (Table 2) for additional analyses because they contained exosome-derived data from 10 patients with primary hepatobiliary tumor-HCC, and 4 patients with benign hepatobiliary tumors. The R-based GEO (Davis et al., 2007) query data package was used for downloading datasets from the GEO database. GSE106452 was not selected for further analysis as the data were generated from HCC cell lines with differing metastatic properties, and no comparison with a normal (non-transformed) cell line was provided.

**Table 2.** List of samples selected from GSE209611 dataset.

| HCC Exosomes |  |  | Benign Exosomes |  |  |
| --- | --- | --- | --- | --- | --- |
| ID | Gender | Age | ID | Gender | Age |
| GSM6380665 | Male | 66 | GSM6380675 | 65 | Female |
| GSM6380666 | Male | 66 | GSM6380676 | 55 | Female |
| GSM6380667 | Female | 68 | GSM6380677 | 76 | Male |
| GSM6380668 | Male | 62 | GSM6380678 | 77 | Female |
| GSM6380669 | Male | 63 |  |  |  |
| GSM6380670 | Male | 71 |  |  |  |
| GSM6380671 | Female | 66 |  |  |  |
| GSM6380672 | Male | 75 |  |  |  |
| GSM6380673 | Male | 71 |  |  |  |
| GSM6380674 | Male | 75 |  |  |  |

To identify differentially expressed miRNAs from the 14 samples in the dataset GSE209611 (Table 2), the miRNA expression data in txt format were preprocessed. We utilized *limma* (Ritchie et al., 2015) to calculate the differential expression (DE) of miRNAs derived from the exosomes of benign and non-benign tumors using R (version 1.4.1717). Significance was determined based on a False Discovery rate (FDR) threshold of < 0.05 and LogFC>1.

#### 2.5.2 Target genes prediction and gene ontology (GO) analysis

microRNAs were analyzed for their potential targets using the MIENTURNET tool (Licursi et al., 2019). To ensure high confidence in the results, FDR<0.05 and a minimum of 4 interactions (each gene should be targeted by at least 4 miRNAs) were set as thresholds for enrichment analyses. The findings were validated with the miRwalk prediction database (Sticht et al., 2018) using a set threshold score of 1. The score was calculated from a random-forest based approach by executing the TarPmiR algorithm (Ding et al., 2016) for miRNA target site predictions. This approach shows the probability of a functional interaction based on the training data. Genes that were identified with both miRwalk and MIENTURNET were used for further analyses.

The gene ontology (GO) platform (Aleksander et al., 2023) was utilized for enrichment analysis and functional annotation to identify the unique biological attributes of the predicted target genes. The data were statistically analyzed with Fisher’s test at FDR <0.05. DIANA-mirPath V3.0 (Vlachos et al., 2015) was used to determine the significantly enriched Kyoto Encyclopedia of Genes and Genomes (KEGG) pathways (FDR < 0.05).

#### 2.5.3 Prognostic evaluation of exosomal miRNAs

We next used the miRNA expression data available in the TCGA database [Liver Hepatocellular Carcinoma (LIHC) data] (Shimada et al., 2019) to evaluate the prediction of prognosis. The expression data was downloaded using the GDC data transfer tool. We normalized the data and then applied log-transformation to ensure comparability and to reduce skewness. After the clinical data was matched with the samples, a total of 362 patients with both miRNA expression data and survival information were selected for further analysis. Clinical data of these patients is shown in Table 3.

**Table 3.** Clinical characteristics of LIHC TCGA cohort examined in the current study.

| Category | Factor | Number |
| --- | --- | --- |
| Project ID | LIHC | 362 |
| Gender | Male | 248 |
|  | Female | 114 |
| Age | >62 | 165 |
|  | ≤62 | 197 |
| Pathological Staging | Unknown | 20 |
|  | Stage I | 173 |
|  | Stage II | 83 |
|  | Stage III | 3 |
|  | Stage IV | 61 |
|  | Stage IIIA | 9 |
|  | Stage IIIB | 9 |
|  | Stage IIIC | 1 |
|  | Stage IVA | 1 |
|  | Stage IVB | 2 |

To identify potential prognostic miRNAs, a univariate Cox regression analysis was carried out for a subgroup of miRNAs (please see Table 5), with overall survival (OS) as the dependent variable and the miRNA expression level as the independent variable at FDR ≤ 0.05. For the construction of a prognostic miRNA signature, we employed multivariate Cox regression analysis using the *survival* package in R to derive multivariate Cox regression coefficients for the subset of miRNAs with positive univariate Cox coefficients. These coefficients were then used to weigh the expression levels of the miRNAs. By combining the weighted expression levels using a risk score calculation method, we obtained a single prognostic score for each patient. Subsequently, the patients were categorized into two groups based on their prognostic scores: high-risk and low-risk. The optimal cutoff point for stratifying patients into these groups was determined using maximally selected rank statistics (MSRS). Kaplan-Meier (KM) survival analysis was conducted to estimate the survival curves for the two groups, and log-rank test was employed to evaluate the statistical significance of the difference between the groups.

#### 2.5.4 Prediction of patient status using a machine learning model

The expressions of miRNAs were correlated with survival data and the miRNAs that were significantly associated with poor prognosis were identified. The expression of these miRNAs in LIHC patients (n=362) in the TCGA cohort was used as input features for a classification model, which was trained to predict the patient’s status (dead or alive) (Table 4).

**Table 4.**
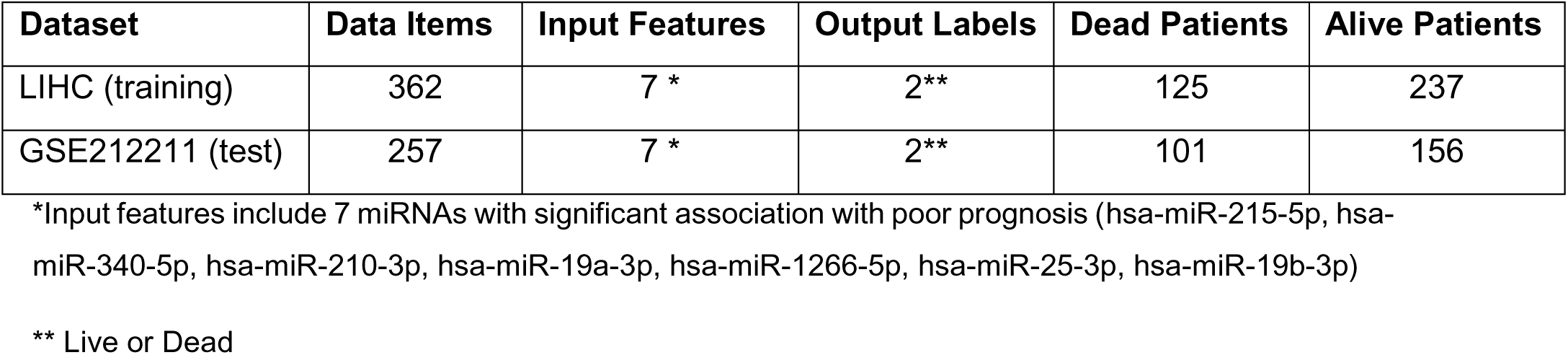
A summary of the datasets, including the distribution of the target variable (dead or alive).

| Dataset | Data Items | Input Features | Output Labels | Dead Patients | Alive Patients |
| --- | --- | --- | --- | --- | --- |
| LIHC (training) | 362 | 7 * | 2** | 125 | 237 |
| GSE212211 (test) | 257 | 7 * | 2** | 101 | 156 |

\*Input features include 7 miRNAs with significant association with poor prognosis (hsa-miR-215-5p, hsa-miR-340-5p, hsa-miR-210-3p, hsa-miR-19a-3p, hsa-miR-1266-5p, hsa-miR-25-3p, hsa-miR-19b-3p)
\*\* Live or Dead

For the training of the classification model, initially, the data was scaled and shifted to have a mean of zero and standard deviation of one. Five-fold cross validation was used to evaluate the model, in order to mitigate overfitting and provide a more reliable estimate of the model’s generalization performance.

As our dataset was imbalanced (the number of alive patients was much higher than the number of dead patients, Table 4), the BalancedRandomForest classifier from the imbalanced-learn package was utilized in Python. This classifier oversamples the minority class (with replacement) during training. To evaluate the performance of the model, the F1-score and AUROC (Area Under the Receiver Operating Characteristic Curve) were determined. The AUROC scores vary between 0 and 1, with values closer to 1.0 indicating higher predictive accuracy, whereas an AUROC of 0.5 would approximately be a random guesser.

The F1-score is a commonly used metric in binary classification and imbalanced learning scenarios. It combines precision and recall scores to provide a single metric of the model’s predictive performance. The ROC curve is another important indicator for imbalanced binary classification. It plots the trade-off between recall (True Positive Rate) and the False Positive Rate (FPR), which measures the rate of negative samples incorrectly classified as positive. The AUROC curve reflects the model’s ability to balance recall and FPR, indicating its overall performance.

To test the performance of our trained model, we utilized a publicly available dataset (GSE212211) which includes 257 individuals with hepatocellular carcinoma (HCC), 41 individuals with cholangiocarcinoma carcinoma (ICC), and two individuals with gall bladder cancer. We focused specifically on the miRNA levels of the 257 HCC patients prior to treatment. Among these patients, 156 were classified as alive, while 101 were classified as dead (Table 4). The details of these patients are shown in Supplementary Table 1. The training dataset was normalized with respect to the mean and standard deviation, and the test dataset was then normalized based on the same mean and variance of the training data. This ensured that there was no leakage from train to test data.

### 2.6 Construction of a Bayesian Network

A Bayesian Network based on domain knowledge was constructed. This made it possible to merge different data sets into one graphical representation, and enabled us to select miRNAs that may have a greater influence on EMT and survival status along with the genes/pathways that they regulate (Figure 1).

**Figure 1.**
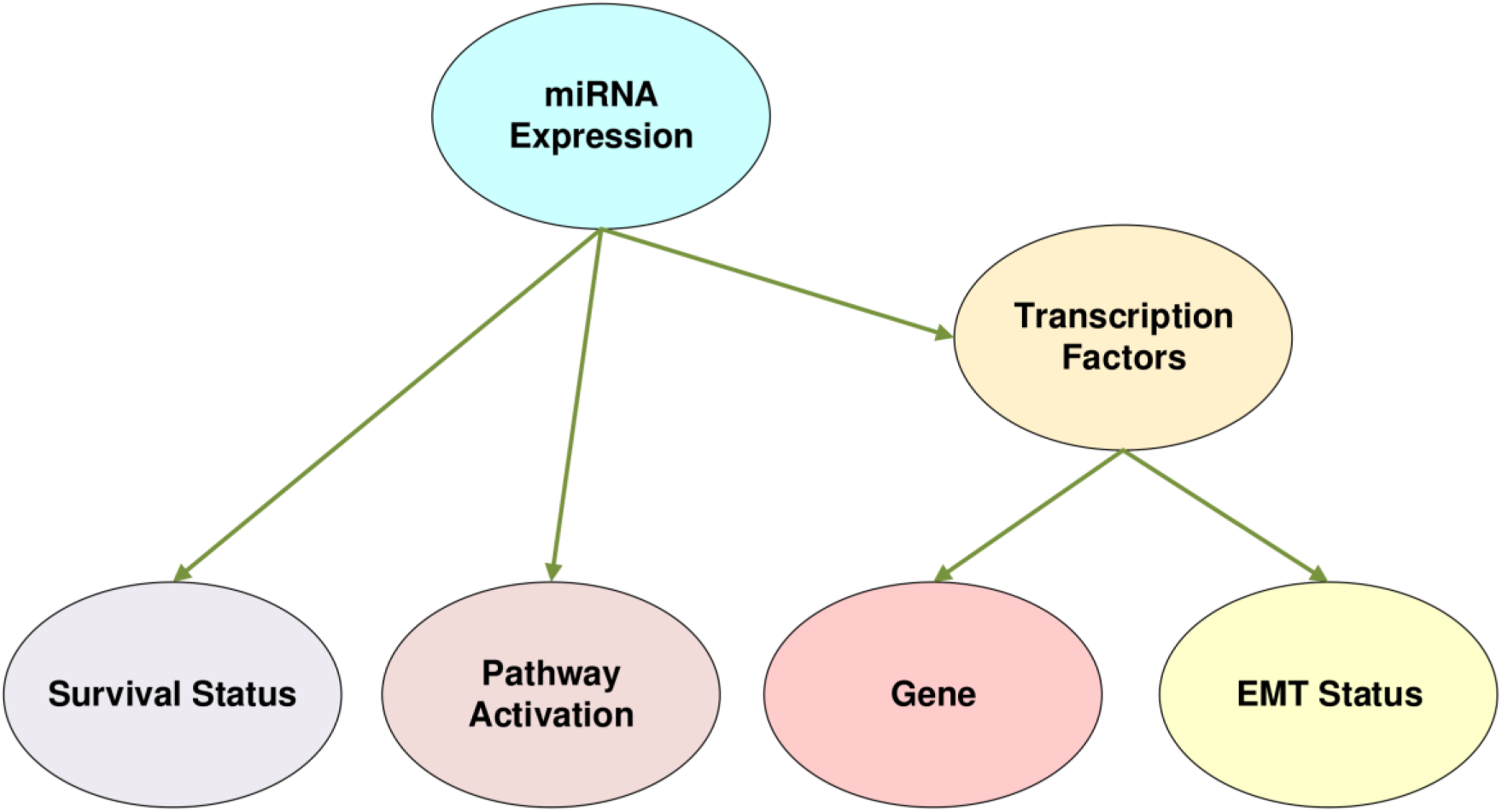
A Bayesian network for the investigation of the post-probability distribution of the miRNAs. EMT: Epithelial to mesenchymal transition. The network was constructed with GeNIe Academic Version 4.0.2423.0

The nodes, the data sets to construct the nodes, the dependencies between the nodes, and the prior probability tables of the nodes were compiled follows (Figure 1):

1. *miRNA node:* The miRNAs that were found to affect patient survival significantly were all assigned equal probability as a prior distribution. This node was chosen as the parent node of all other nodes since the miRNAs are presumed to causally influence the other entities.
2. *Survival status node*: The survival of patients associated with the expression of miRNAs were assigned binary codes (1-0) as prior probabilities. This node was conditionally linked to the miRNA node only, because conditional probabilities were generated from the survival data used in this study.
3. *Pathway activation node:* This node was created by determining the pathways that were regulated by the miRNAs associated with survival. The pathways were identified from MiRPathDB version 2.0 (Kehl et al., 2020) and the most significant pathways were grouped under a general heading (Supplementary Table 2, Tab: mirna pathways-miRPathDB2). This node was conditionally connected to the miRNA node, and the pathway status was assigned with binary (1-0) codes for prior probability tables based on whether they were associated with the miRNA or not.
4. *Transcription Factors node*: This node was created to evaluate the relationship between the miRNAs and the target genes that were significantly enriched by a GO analysis. For this node, we used the TransmiR database (Tong et al., 2019), which provides a list of transcription factors (TF) that are regulated by microRNAs. The TFs regulated by the miRNAs that were significantly associated with survival were retrieved from this database (Supplementary Table 2, Tab: TransmirDB_mir_TF_mirnas) and assigned to the miRNAs as a binary code (1-0). This node was defined as the Child node of the miRNA node to represent the causal relationship between microRNAs and TFs.
5. *EMT node*: To analyze whether the TFs in the TFs node were related to the EMT process, all human EMT-related TFs (Supplementary Table 2, Tab: all human EMT genes -dbemt.bioi) that were identified from Epithelial-Mesenchymal Transition gene database (Zhao et al., 2019) were compared with TFs in the TFs node. The EMT-related TFs and unrelated TFs were categorized as binary codes (1-0) and added to the network. This node was defined as a Child node of TFs since the TFs could affect EMT.
6. *GENE node*: In this node, genes from enrichment analysis using GO were connected to the TFs node with binary codes (0-1) to represent their relation. The data for the node were created from the TF2DNA Database (https://www.fiserlab.org/tf2dna_db/), by selecting *Homo sapiens*, TF2DNA [Computational] as source, and p-value=0.0001. From the results, promoter regions of the enriched genes were selected and connected with TFs nodes as Child nodes (Supplementary Table 2, Tab: TFs related to 4 genes). An additional row called *not-related* was added to this node to represent the genes other than the significantly enriched genes that can also be affected by the miRNAs. The binary codes for this row were assigned as 1 if there was no relation between TFs and the enriched genes, and 0 if there was at least one relation between the TFs and the enriched genes.

During the creation of the data sets as Excel files, all duplicated values were removed. All binary codes that were used for the creation of the prior probability tables of the network were generated in R version 2022.12.0 by using the data sets mentioned. The binary codes were fed into the Bayesian network and all values were normalized to create the prior probability tables. All binary codes for the construction of the Bayesian network are provided in Supplementary Table 2 (Tab: binary code to feed network), and the dataset from the Bayesian network after the creation of the network that was used for the validation step is shown in Supplementary Table 2 (Tab: data for validation). Sensitivity analyses were carried out before the validation step for different target nodes on the network. With *Validation* option in *Learning* menu in the GeNIe software, desired nodes were selected to operate validation and since the data sets for creation of the network and testing of the model are the same, and included a relatively small number of samples; five k-fold cross-validation was used to train the model and validate the results.

### 2.7 Identification of strong miRNA predictors

To identify the miRNAs that can strongly predict survival, the accuracy result of each miRNA was considered individually. Among the miRNAs associated with survival, those that had the highest accuracy scores after validation were selected to obtain posterior probabilities. The posterior probabilities of the remaining miRNAs were ignored. After setting conditions on the nodes of survival status (SS), EMT (EMT) and gene (G), posterior probabilities of the miRNAs (M), TFs (TF) and pathway status (PS) were created.

### 2.8 Statistical analyses

The statistical method used for each analysis is described under the relevant image in the figure legend or in the methodology.

## 3. Results

### 3.1 Identification of differentially expressed miRNAs and their target genes

Exosomes isolated from HuH7 cells overexpressing Slug showed up-regulation of fifty-four exosomal miRNAs compared to the control cells (Supplementary Table 3). We randomly selected three miRNAs from these fifty-four upregulated miRNAs and validated their upregulation in HuH7 cells undergoing p-EMT (via the overexpression of Slug) compared to the empty vector control (Supplementary Figure 1) and could validate the upregulation of all three miRNAs. We next examined publicly available datasets that compared the miRNA expression profile from exosomes collected from non-benign HCC tumors versus exosomes collected from benign tumors (GSE209611). Among the top 250 differentially expressed miRNAs, 56 were found to be significantly up-regulated in non-benign exosomes (Supplementary Table 4), while 194 miRNAs were down-regulated. Of the upregulated miRNAs, 13 were shared with the miRNAs that were upregulated in exosomes from HuH7 cells expressing Slug, suggesting that these miRNAs may be important for EMT and/or invasion (Table 5).

**Table 5:** miRNAs that were upregulated in exosomes from Slug overexpressing HuH7 cells and in exosomes isolated from HCC tumors.

| miRNA | miRBase Accession |
| --- | --- |
| hsa-miR-375 | MIMAT0000783 |
| hsa-miR-215-5p | MIMAT0000272 |
| hsa-miR-340-5p | MIMAT0004692 |
| hsa-miR-210-3p | MIMAT0000267 |
| hsa-miR-19a-3p | MIMAT0000073 |
| hsa-miR-1266-5p | MIMAT0005920 |
| hsa-miR-25-3p | MIMAT0000081 |
| hsa-miR-1303 | MIMAT0006370 |
| hsa-miR-19b-3p | MIMAT0000074 |
| hsa-miR-296-3p | MIMAT0004679 |
| hsa-miR-218-5p | MIMAT0000275 |
| hsa-miR-101-3p | MIMAT0000099 |
| hsa-miR-885-5p | MIMAT0004947 |

To identify whether any of these miRNAs were of prognostic significance in human HCC, we used the pipeline described in Scheme 1.

**Scheme 1:**
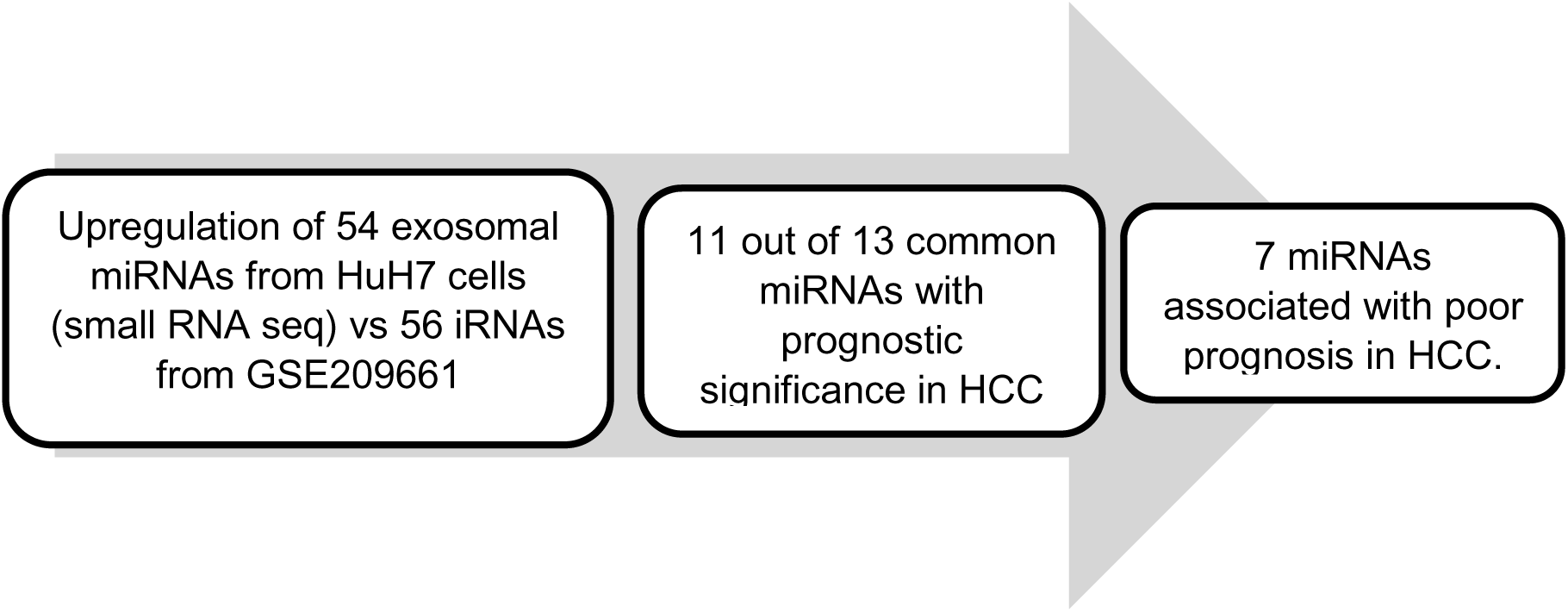
Pipeline for the identification of prognostically significant exosomal miRNAs.

Upregulation of 54 exosomal miRNAs from HuH7 cells (small RNA seq) vs 56 iRNAs from GSE209661 11 out of 13 common miRNAs with prognostic significance in HCC 7 miRNAs associated with poor prognosis in HCC.

### 3.2 Functional annotations and enrichment analysis of the 13 miRNAs

The 13 common miRNAs that were upregulated in exosomes isolated from Slug overexpressing HuH7 cells as well as exosomes isolated from human HCC (GSE209611) (Table 5) were examined further for their targets. For this, we used the MIENTURNET tool with an FDR threshold of ≤0.05 and a minimum interaction requirement of 4 miRNAs per gene. This led to the identification of target 81 genes (Supplementary Table 5) that were also validated independently with the miRwalk database. Four genes, *BCL2L11, DICER1, MLEC,* and *SLC7A11*, exhibited the highest number of interactions, indicating that they are potentially targeted by the highest number of miRNAs in our list (Supplementary Table 5).

We next carried out GO functional annotation analysis for all 81 targets of the 13 miRNAs (Figure 2) and observed a general enrichment of terms associated with cytoskeletal organization, miRNA processing, mesenchymal cell proliferation, protein processing in the ER and signal transduction.

**Figure 2:**
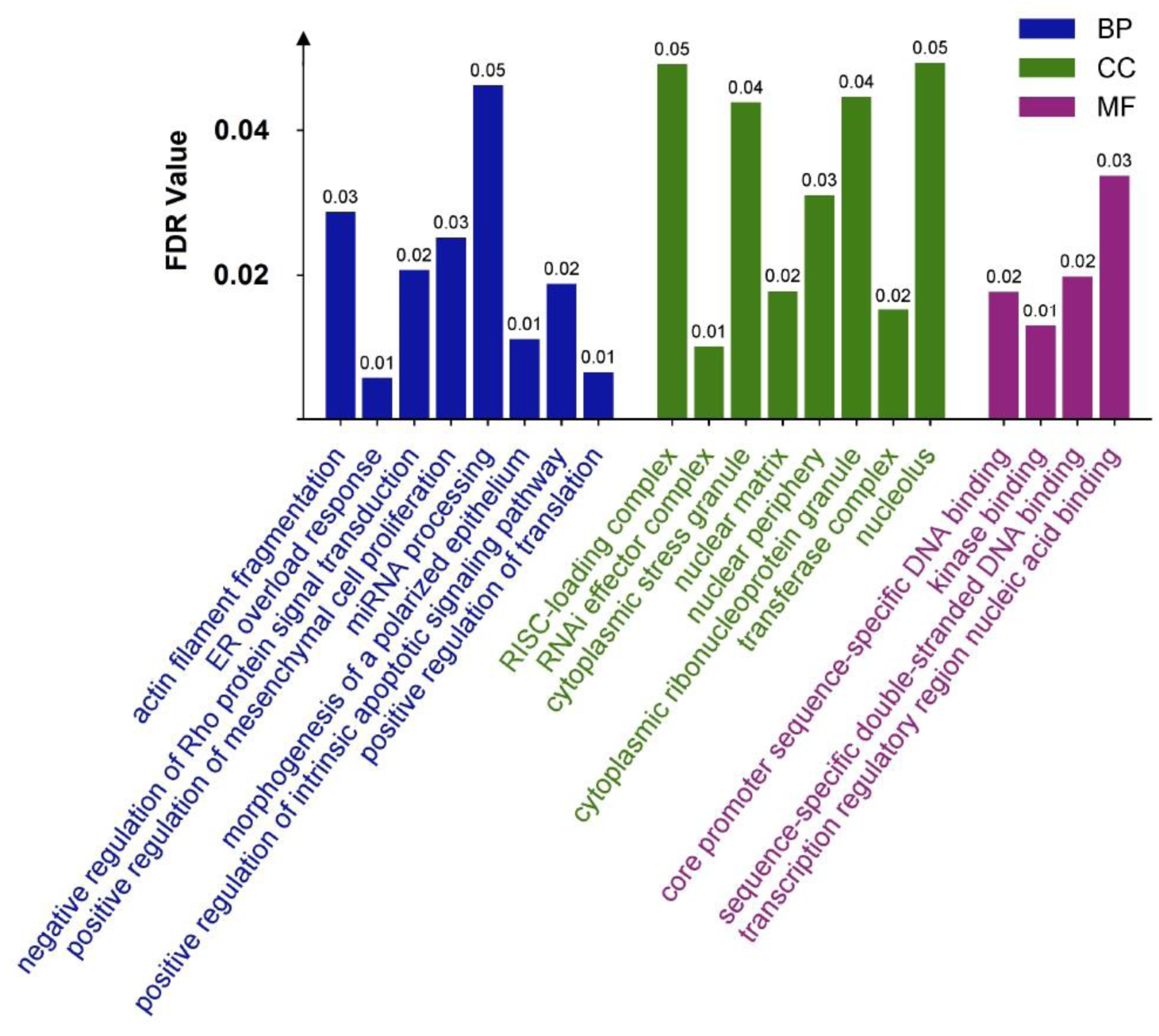
Gene ontology (GO) functional annotation analysis for the targets of the 13 miRNAs terms (P < 0.05). Terms association with Biological Processes (BP), Cellular Compartment (CC) and Molecular Functions (MF) are shown.

Next, we carried out a KEGG pathway enrichment analysis which indicated significant enrichment of cell-cell junctions, extracellular matrix and cytoskeleton, signal transduction (such as via TGFβ, FOXO and p53), transcriptional deregulation in cancer and protein processing in the ER (Figure 3).

**Figure 3:**
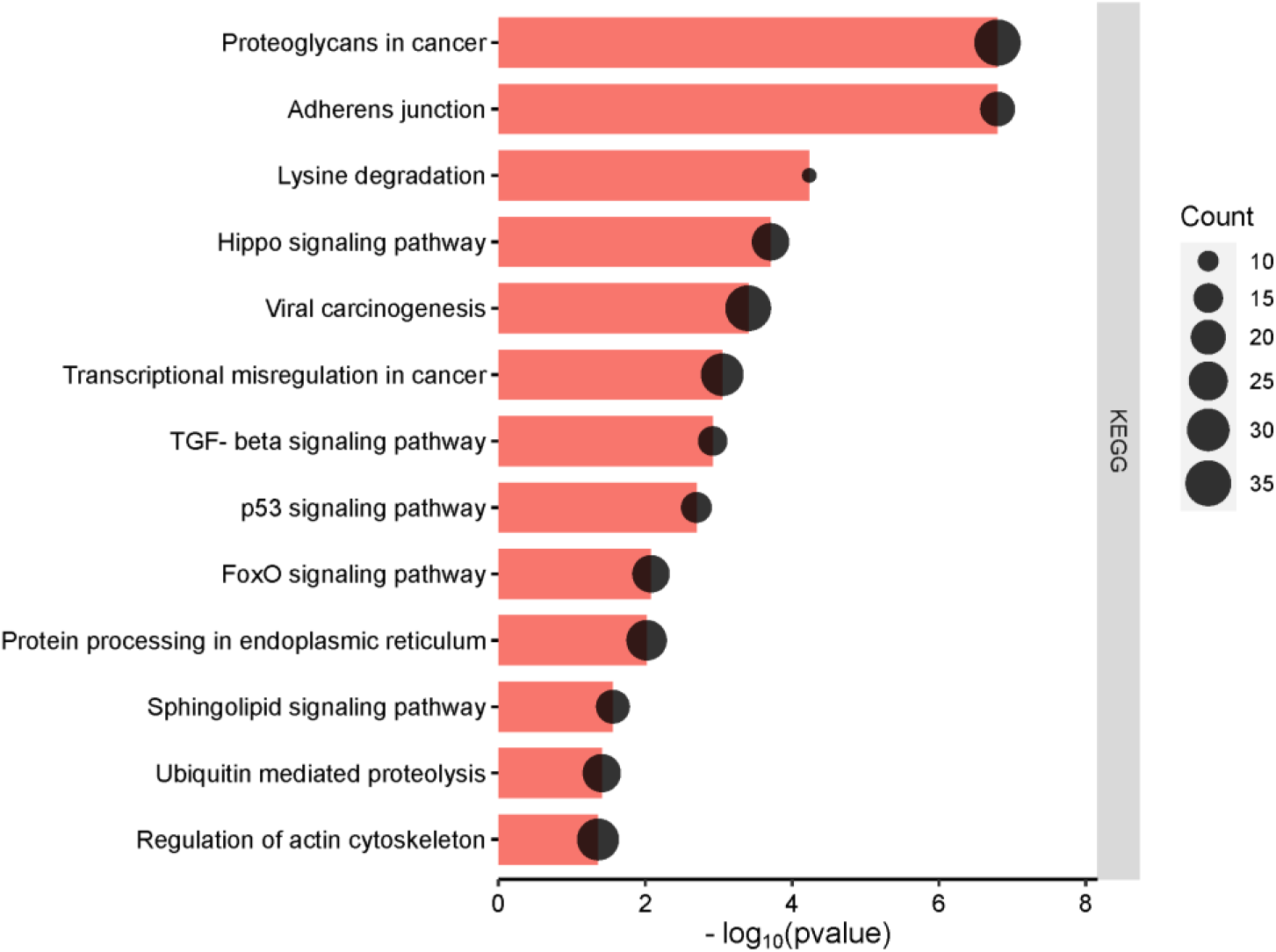
KEGG pathway enrichment analysis using the significant (p<0.05) targets of the 13 miRNAs. Count refers to the number of miRNAs that are targeting genes in that pathway.

### 3.3 Survival analysis using TCGA LIHC data

We next evaluated whether the 13 miRNAs were of prognostic significance in HCC. For this, we analyzed the overall survival (OS) of the Liver Hepatocellular Carcinoma (LIHC) cohort based on the expression of each of these miRNAs. Using univariate Cox regression analysis, 11 miRNAs exhibited a statistically significant association with patient survival (p < 0.05) (Table 6). Of these, the high expression of seven miRNAs, namely, hsa-miR-215-5p, hsa-miR-340-5p, hsa-miR-210-3p, hsa-miR-19a-3p, hsa-miR-1266-5p, hsa-miR-25-3p, and hsa-miR-19b-3p was significantly associated with reduced OS. Four miRNAs, namely, hsa-miR-375, hsa-miR-218-5p, hsa-miR-101-3p, and hsa-miR-885-5p presented negative Cox coefficients, indicating that higher expression levels were associated with a decreased risk of death, suggesting a potentially beneficial impact on OS.

**Table 6.**
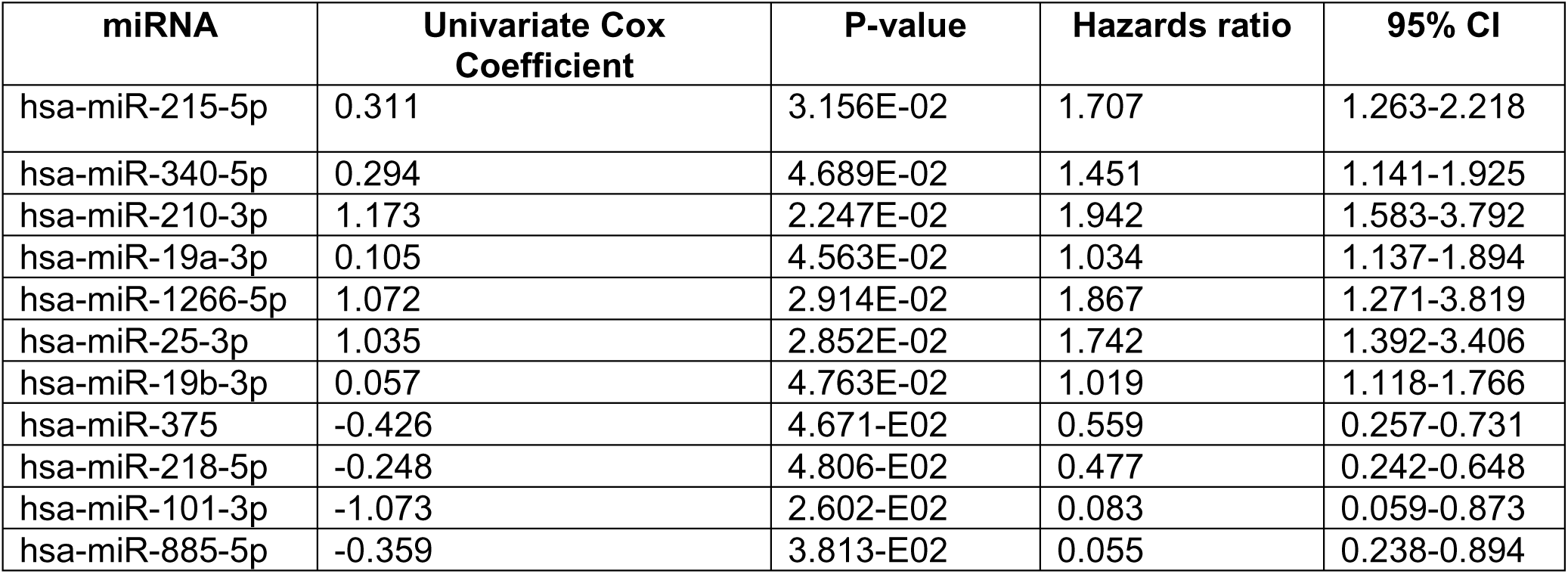
Univariate Cox regression analysis of the 11 miRNAs that showed prognostic significance in HCC.

| miRNA | Univariate Cox Coefficient | P-value | Hazards ratio | 95% CI |
| --- | --- | --- | --- | --- |
| hsa-miR-215-5p | 0.311 | 3.156E-02 | 1.707 | 1.263-2.218 |
| hsa-miR-340-5p | 0.294 | 4.689E-02 | 1.451 | 1.141-1.925 |
| hsa-miR-210-3p | 1.173 | 2.247E-02 | 1.942 | 1.583-3.792 |
| hsa-miR-19a-3p | 0.105 | 4.563E-02 | 1.034 | 1.137-1.894 |
| hsa-miR-1266-5p | 1.072 | 2.914E-02 | 1.867 | 1.271-3.819 |
| hsa-miR-25-3p | 1.035 | 2.852E-02 | 1.742 | 1.392-3.406 |
| hsa-miR-19b-3p | 0.057 | 4.763E-02 | 1.019 | 1.118-1.766 |
| hsa-miR-375 | -0.426 | 4.671-E02 | 0.559 | 0.257-0.731 |
| hsa-miR-218-5p | -0.248 | 4.806-E02 | 0.477 | 0.242-0.648 |
| hsa-miR-101-3p | -1.073 | 2.602-E02 | 0.083 | 0.059-0.873 |
| hsa-miR-885-5p | -0.359 | 3.813-E02 | 0.055 | 0.238-0.894 |

The 7 miRNAs with positive univariate Cox coefficient (Table 6) were used to construct a prognostic miRNA signature. We selected these miRNAs as their expression was associated with poor OS and were more likely to reflect the occurrence of p-EMT. The Cox regression coefficients derived from the multivariate Cox regression analysis were used to weigh the expression levels of each miRNA.

The weighted expression levels used to obtain a single prognostic score for each patient and the patients were stratified into two groups: a high-risk group (n = 150) and a low-risk group (n = 120). The optimal cutoff point for stratification was determined by using a value that maximized the separation of the survival curves. The Kaplan-Meier survival analysis of hsa-miR-215-5p, hsa-miR-340-5p, hsa-miR-210-3p, hsa-miR-19a-3p, hsa-miR-1266-5p, hsa-miR-25-3p, and hsa-miR-19b-3p are shown in Figure 4.

**Figure 4.**
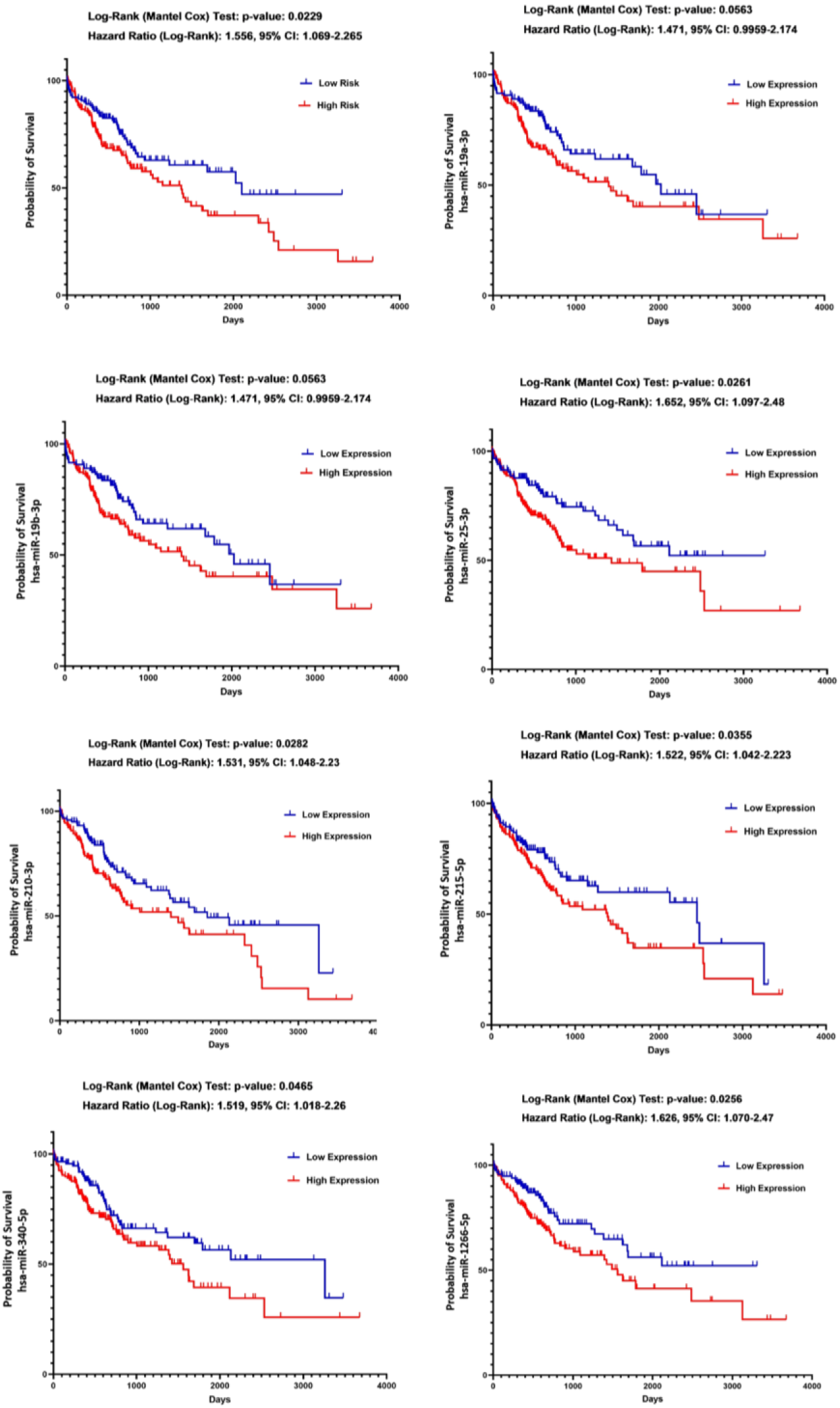
Kaplan-Meier (KM) survival plots for overall survival (OS). (A) KM plots for the expression of all seven miRNAs (hsa-miR-215-5p, hsa-miR-340-5p, hsa-miR-210-3p, hsa-miR-19a-3p, hsa-miR-1266-5p, hsa-miR-25-3p, and hsa-miR-19b-3p). (B) KM plot for hsa-miR-215-5p (C) KM plot for hsa-miR-340-5p (D) KM plot for hsa-miR-210-3p (E) KM plot for hsa-miR-19a-3p (F) KM plot for hsa-miR-1266-5p (G) KM plot for hsa-miR-25-3p (H) KM plot for hsa-miR-19b-3p. Patients with high expression levels of these miRNAs exhibited significantly worse prognosis compared to those with low expression levels.

The slight disparity observed between the univariate Cox coefficient (Table 6) and the Log-Rank test results (Figure 4) can be attributed to the differences in the data utilized for these analyses. The univariate Cox coefficient was calculated using the complete set of patients (362 patients), while the Log-Rank test was performed after stratifying the patients into high-risk and low-risk groups, with a subset of 270 patients selected for the test. This deliberate stratification allowed for a more refined evaluation of patient separation based on the risk score, yielding a more informative outcome.

### 3.4 Generation of a model for prognostication of HCC patients based on exosomal miRNA expression using machine learning

The expression of the seven miRNAs associated with worse prognosis and the vital status available from TCGA were next used with machine learning to examine whether a signature could be generated that can predict survival of HCC patients. Since our dataset is imbalanced (i.e., the number of patients that were dead or alive were not equal), accuracy was not an informative metric. To evaluate the performance of the model, the F1-score and AUROC were calculated for each fold, as well as their average. The average AUROC score and F1-score for the 5-fold cross validation were 0.6848 and 0.6881, respectively.

To test our trained model, we used the test dataset (GSE212211) and observed the AUROC and F1 scores of 0.6859 and 0.7320, respectively (Figure 5). These scores indicate that in GSE212211, patients with higher expression of the seven specific miRNAs (hsa-miR-215-5p, hsa-miR-340-5p, hsa-miR-210-3p, hsa-miR-19a-3p, hsa-miR-1266-5p, hsa-miR-25-3p, hsa-miR-19b-3p) had a higher probability of being deceased, while most of the patients who were alive exhibited lower expression of these miRNAs, consistent with the TCGA data set.

**Figure 5.**
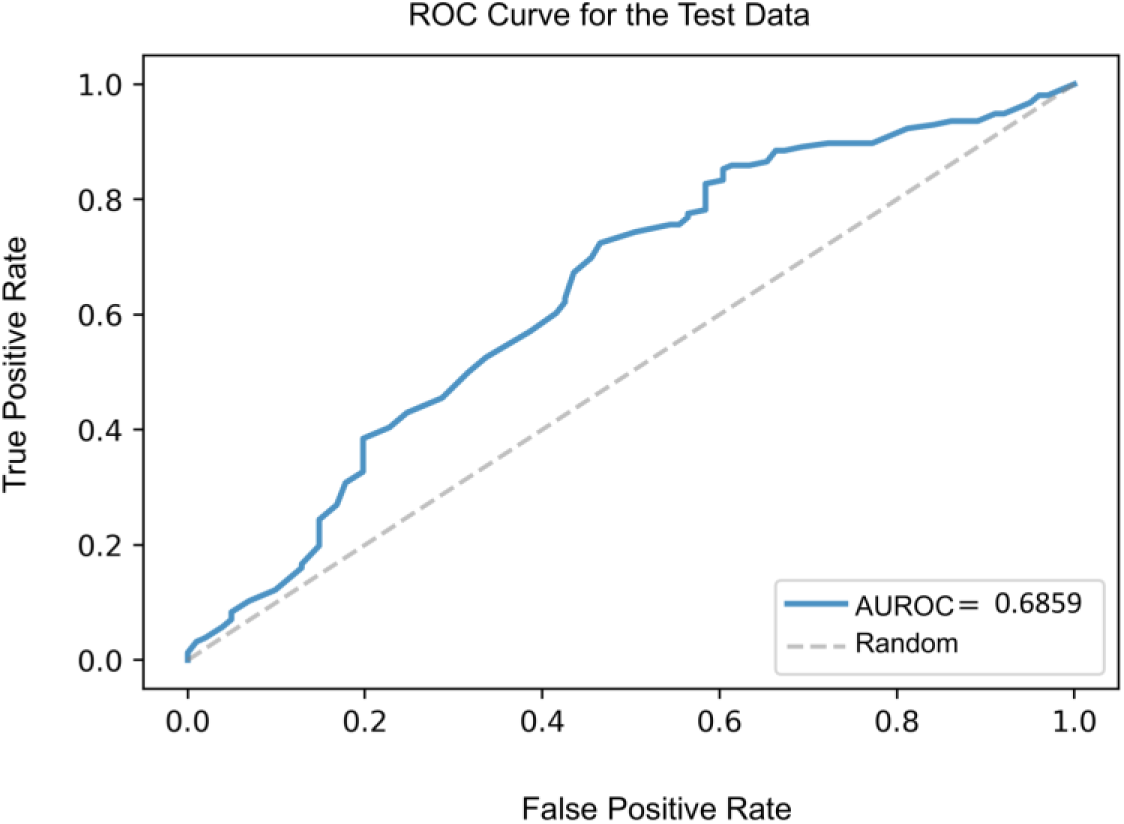
Receiver operating characteristic (ROC) curve for test data (GSE212211) and AUROC showing the performance of the miRNA signature.

### 3.5 Bayesian network to predict the function of specific miRNAs

We next examined which miRNAs were the most effective in inducing p-EMT phenotype and poor prognosis by constructing a Bayesian Network. The nodes that were used for this network are described in the Methods. For Pathway Activation, since the GO terms in the MiRPathDB output included ‘mitotic cell cycle, cell cycle, and chromosome segregation’, the related row in the probability table of the node was given a general name ’cell cycle’. The node includes the following general pathways as rows: Transcription-related, cell cycle-related, neural processes-related, metabolic activities-related, and morphology related (Figure 6). The “EMT” node and “Pathway Activation” nodes were excluded for validation to avoid overfitting of these nodes and the survival status node was excluded since the survival data is fixed.

**Figure 6:**
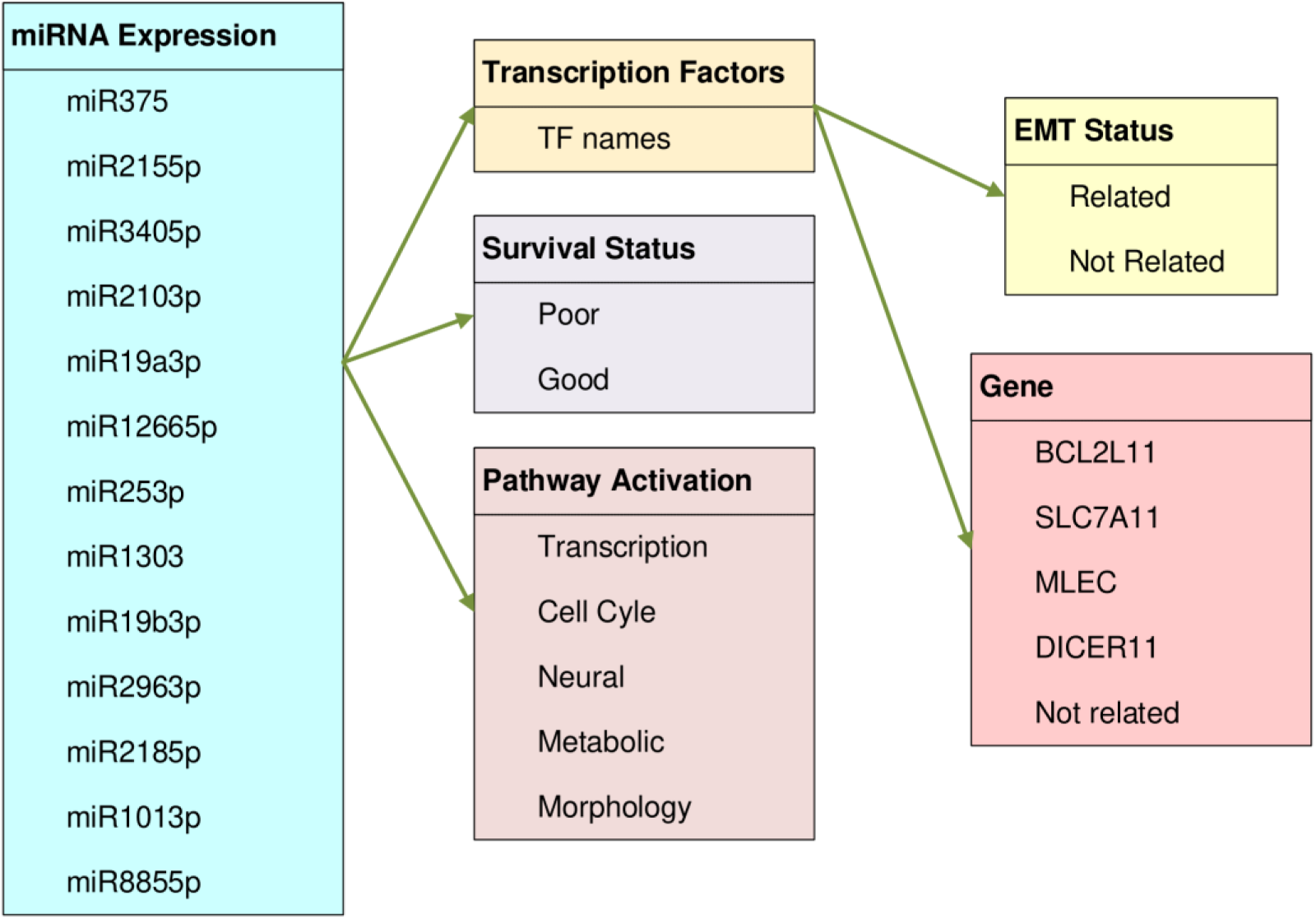
Parent and Child nodes in the Bayesian network. The miRNA node was the Parent node, while the Transcription Factors, Survival Status and Pathway Activation were the Child nodes of the miRNA node. EMT status and Gene are the Child nodes of Transcription Factors. The names of the TFs node were not included in the figure as their number was too high. The network was constructed with GeNIe Academic Version 4.0.2423.0.

Six of the 13 miRNAs had higher validation scores compared to the others. The validation scores are as follows: miR375 = 0.67, miR215-5p = 0.7, miR210-3p = 0.33, miR1266-5p = 0.90, miR19b-3p = 0.85, and miR218-5p = 0.86. Four of these six miRNAs were associated with worse prognosis, while two were associated with favorable prognosis (Table 6). We next examined these validated microRNAs (M) and computed posterior probabilities for their associations to better understand the regulatory network of each of the four significantly enriched genes (*BCL2L11*, *DICER1*, *MLEC*, and *SLC7A11*), poor survival status and EMT-relatedness. For the calculation of posterior probabilities (Figure 7), evidences (E) were selected from the nodes of Survival Status (SS), EMT status and Gene (G) and posterior probabilities of the nodes M, TF and PS were investigated. Since,

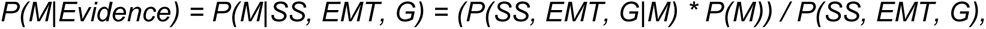

we set evidence of G for each of the 4 genes for each iteration *(G=BCL2L11, G=DICER1, G=MLEC, and G=SLC7A11)* and *SS = poor*, *EMT = related* for all the cases, where *P(poor, related, BCL2L11|M)* is the joint probability of *SS = poor, EMT = related,* and *G = BCL2L11* given *M: P(M)* is the prior probability of M being true. Then, for the selected gene (in this case, *G = BCL2L11*);

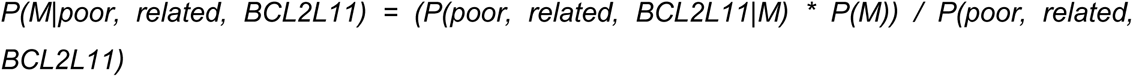

was calculated to represent the probability of node *M being true (M = 1)* given the evidenc*e SS = poor, EMT = related,* and *G = BCL2L11.* All equations were relevant for posterior probabilities of TF and PS, by changing M to TF and to PS.

**Figure 7:**
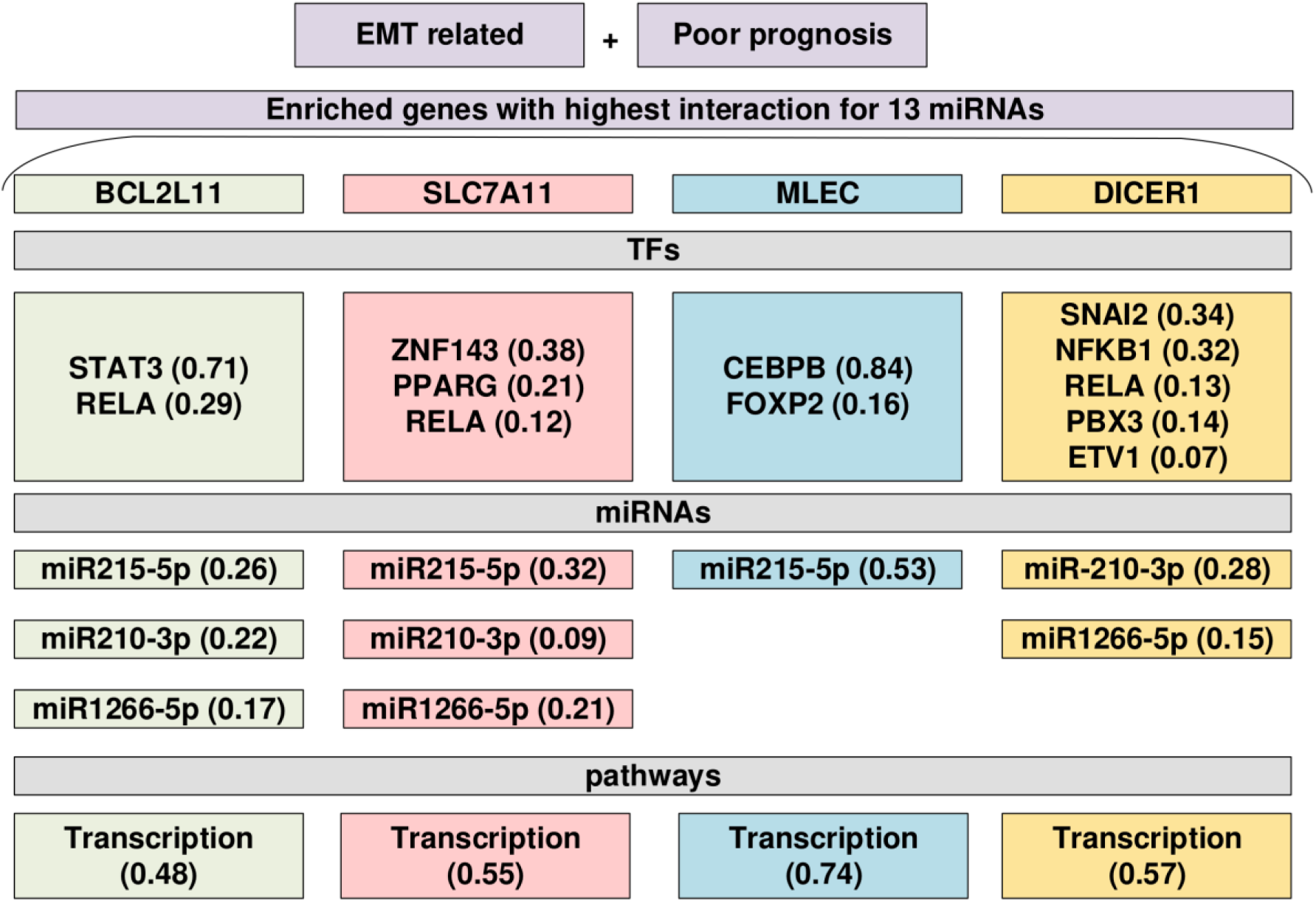
Posterior probability scores (PPS) using Bayesian Inference networks for miRNA, Pathway and Transcription Factor (TF) for the four enriched genes (BCL2L11, SLC7A11, MLEC and DICER1).

Out of the six miRNAs with high validation scores, three were found to be more related to the four genes in the context of EMT and poor prognosis. All three of these miRNAs were found to be associated with poor prognosis (Table 7). For *BCL2L11,* miR215-5p displayed a posterior probability of 0.26, miR210-3p displayed a posterior probability of 0.22, and miR1266-5p displayed a posterior probability of 0.17 among the validated microRNAs (Figure 7).

The impact of various pathways for the settled evidence was also investigated for *BCL2L11* (*P(PS|poor, related, BCL2L11)*). The most significant pathway was found to be the Transcription pathway, which outperformed other pathways with a posterior probability of 0.48 (PS). We also evaluated the role of transcription factors (TFs) in the evidence of *BCL2L11* and among the 263 TFs (*P(TF|poor, related, BCL2L11)*), STAT3 had the highest posterior probability of 0.71, while RELA showed a posterior probability of 0.29.

The same analyses were repeated for SLC7A11, MLEC and DICER1 and posterior probabilities were calculated for the transcription factors and miRNAs (Figure 7). Of note, for all genes, the transcription pathway outperformed all other pathways, suggesting that the p-EMT phenotype was mainly driven by transcription.

## 4. Discussion

Tumors are highly heterogenous; therefore, diagnosis, prevention, therapy and follow-up need to be tailored for individual patients. Partial EMT (p-EMT) was suggested as a poor prognostic factor in hepatocellular carcinoma (HCC) (Lei et al., 2021). Moreover, circulating tumor cells with a p-EMT phenotype were associated with advanced clinical stage, metastasis and higher alpha fetoprotein (AFP) levels in HCC patients (Chen et al., 2017). In this study, we identified fifty-four upregulated exosomal miRNAs in Huh7 cells undergoing partial EMT (Karaosmanoğlu et al., 2018). Exosomes are secreted by numerous cell types (epithelial, immune etc.) into the extracellular space and can be found in various bodily fluids. Therefore, exosomes have gained high significance in recent years as potential biomarkers that can predict disease outcomes through non-invasive liquid biopsy (Liu et al., 2023). To evaluate the relevance of these miRNAs to HCC, we evaluated the publicly available dataset GSE209611 and identified 13 common miRNAs that were also upregulated in exosomes collected from HCC tumors, suggesting that these miRNAs may be relevant to disease.

The 13 common miRNAs had a total of 81 mRNA targets, which were functional in a number of different cellular processes such as extracellular matrix, cytoskeletal organization, cell-cell junction, mesenchymal cell proliferation, signal transduction, protein processing and translation. Many of these processes are highly relevant to EMT (Scott et al., 2019). Further evaluation of the 81 targets indicated 4 that were targeted by several (>4) of the 13 common miRNAs. *BCL2L11* (BIM) is a member of the BCL2 family of proteins that induces apoptosis (Sionov et al., 2015). The gene that had the strongest negative correlation with the expression of *BCL2L11* was reported to be the mesenchymal marker *VIM* among 19,000 genes in 857 solid tumor cell lines (Song et al., 2018). In HCC, siRNA mediated suppression of *SNAI1* expression was shown to enhance apoptosis by increasing the expression of *BCL2L11* (Franco et al., 2010). These data suggest that a miRNA mediated suppression in the expression of *BCL2L11* can lead to decreased apoptosis, enhanced cell survival and EMT.

DICER is known to be a negative regulator of EMT. In breast cancer cells, miR-103/107 was reported to directly target DICER1, leading to an overall decrease in the synthesis of miRNAs. Specifically, miR-200, which is known to suppress the mesenchymal marker ZEB1, was lost in Dicer1 suppressed cells, leading to a remarkable increase in EMT (Martello et al., 2010). In HCC, hypoxia was reported to lead to a loss of Dicer, which favored the hypoxia induced induction of EMT (Ibrahim et al., 2017).

Malectin (MLEC) is an ER resident protein that was shown to be induced during ER stress to inhibit the secretion of glycoproteins by decreasing efficient protein de-glucosylation (Galli et al., 2011). Although a specific role of Malectin in the development of EMT in HCC has not been reported to date, altered expression of cell surface glycans has been reported in HCC cell lines undergoing growth factor mediated EMT (Li et al., 2013).

SLC7A11, also known as xCT, is an amino acid antiporter that promotes cystine uptake in exchange of glutamate. The cystine is then used for glutathione biosynthesis for mitigation of oxidative stress and death from ferroptosis (Koppula et al., 2018). During metastasis, cancer cells are more susceptible to ferroptosis due to various mechanisms such as enhanced synthesis of ROS and iron accumulation. Cells undergoing metastasis have also developed various means to inhibit ferroptosis, including increased synthesis of glutathione via increased cystine uptake through SLC7A11 (Jiang et al., 2023). Non-coding RNA (including miRNA) mediated regulation of SLC7A11 has been reported in many different tumor types (Jiang et al., 2023). In HCC, the noncoding RNA Circ0097009 was reported to sponge miR-1261 thereby abrogating miR-1261 mediated suppression of SLC7A11 and enhancing cell survival and metastasis (Lyu et al., 2021).

We next interrogated whether the 13 miRNAs were of prognostic significance. A univariate Cox regression analysis indicated that 11/13 miRNAs were significantly associated with prognosis (OS) in HCC; 7 with worse OS and 4 with favorable OS. We next focused on the 7 miRNAs associated with worse prognosis and observed that expression of these 7 miRNAs in the LIHC cohort of TCGA could predict worse prognosis with AUROC and F1 scores of 0.6848 and 0.6881, respectively, and validated in an independent microarray dataset with AUROC and F1 scores of 0.6859 and 0.7320, respectively.

Most of the 7 miRNAs in the prognostic signature have been reported to be associated with oncogenic functions in HCC. miR-215-5p was reported to be higher in serum exosomes isolated from HCC patients and was associated with poor disease-free survival (DFS) (Cho et al., 2020). A high level of miR-210-3p was also identified in serum exosomes of HCC patients and was associated with enhanced angiogenesis through the suppression of STAT3 and SMAD4 in endothelial cells (Lin et al., 2018). miR-19a-3p was reported to be upregulated in HCC tissues and could enhance metastasis and resistance to sorafenib by targeting phosphatase and tensin homolog (PTEN)/AKT signaling (Jiang et al., 2018). miR-19b-39 was reported to target RNA binding motif single stranded interacting protein 1 (RBMS1) in HCC; the latter protein was found to inhibit the expression of glutathione peroxidase 4 (GPX4) and facilitated cell death by ferroptosis. Low expression of RBMS1 was found to be an independent prognostic factor for worse OS and RFS in HCC (Zhai et al., 2023). miR-1266 was also found to be expressed significantly more in HCC tissues compared to normal and was associated with worse OS. Overexpression of miR-1266 in HCC cell lines was associated with enhanced proliferation, migration and invasion (HUANG et al., 2021). miR-25-3p was reported to be expressed more in HCC compared to normal tissues and high expression was associated with worse prognosis. Mechanistically, miR-25-3p was shown to target the tumor suppressor protein F-box and WD repeat domain containing 7 (FBXW7) in order to activate autophagy and enhance resistance to sorafenib (X. Feng et al., 2022). miR-340-5p is the only miRNA from the signature that has been associated with context dependent tumor suppressive or oncogenic functions. The expression of miR-340-5p was found to be lower in HCC tissues a cohort of Chinese patients compared to matched normal tissues. miR-340-5p could decrease proliferation, invasion and migration and enhance apoptosis in HCC cell lines (Wang et al., 2017). However, the expression of miR-340-5p was reported to be enhanced in thyroid carcinoma and gastric carcinoma (Huang et al., 2021). Using the TCGA LIHC data, we observed that high expression of miR-340-5p was associated with a significantly worse OS (Hazards Ratio 1.451, 95% CI 1.141-1.925). This miRNA was also retained in the signature that we developed with a machine learning algorithm in both training and test datasets. It is possible that the expression of this specific miRNA is affected by characteristics of the cohort, even when examined for the same cancer.

Since the AUROC for the prediction of prognosis was below 0.8 and therefore can be considered to be modest (de Hond et al., 2022), we next carried out probabilistic modeling using Bayesian inference networks that can be used to model uncertainty, reasoning over imprecise data sets and infer dependencies between parameters. Of the 13 miRNAs that we initially identified as relevant to HCC, six had high validation scores and were used for the model. Three of the six validated miRNAs (miR 215-5p, miR 210-3p and miR-1266-5p), all associated with worse prognosis, were found to be strongly related to the enriched genes in the context of EMT and poor prognosis.

We computed posterior probabilities to evaluate the association of these miRNAs with the four significantly enriched genes (*BCL2L11*, *DICER1*, *MLEC*, and *SLC7A11*), poor survival status and EMT-relatedness. Remarkably, among the various pathways that could be impacted by the miRNAs and their four primary targets were transcription factors (TF). This finding suggests that the p-EMT phenotype observed was very likely to be driven by transcription. Transcription factors such as Snail, Slug, Twist and Zeb are well known to orchestrate many different aspects of EMT and are considered to be the core EMT-TFs, and are activated during development, wound healing or in diseased states such as cancer (Debnath et al., 2022). All of these transcription factors suppress the expression of E-cadherin (CDH1), thereby compromising cell-cell adherence junctions and enhancing cell motility. The EMT-TFs have multiple additive effects as they can transcriptionally regulate each other’s expression. Moreover, these TFs can functionally cooperate on the expression and activity of genes that they co-target (Lee and Kong, 2016). Tens of additional TFs have been identified that primarily regulate cancer associated EMT, such as proteins of the KLF, SOX, RUNX and forkhead box (FOX) families. These TFs function by targeting the core EMT-TFs (Debnath et al., 2022).

miRNAs are also well known to regulate the expression of EMT-TFs, primarily decreasing their expression and thereby inhibiting EMT. The best described example is the feedback loop between ZEB1 and the miR-200 family that mediates cell differentiation (J. Feng et al., 2022). Of note, EMT-TFs can bind to repressor elements in the promoters of the EMT related miRNAs and thereby inhibit their expression.

Among the 263 TFs evaluated in the model where BCL2L11 was the “gene”, the TF STAT3 had the highest posterior probability of 0.71, while the model where MLEC was the “gene” the TF CEBPB had the highest posterior probability of 0.84. STAT3 is widely known to be associated with EMT in many tumor types including HCC (Sadrkhanloo et al., 2022). Knockdown of STAT3 in HCC cell lines was shown to inhibit angiogenesis via a decrease in vascular endothelial growth factor, apoptosis via decreased expression of survivin, and motility via decreased expression of matrix metalloproteinases 2 and 9 (Li et al., 2006). miRNAs such as miR-200 and Let-7 were reported to be STAT3 effectors in the regulation of EMT (Guo et al., 2013).

C/EBPβ, encoded by the gene *CEBPB* has been reported to be both negatively and positively associated with the development of EMT. C/EBPβ was shown to transcriptionally upregulate miR-29a-3p; the latter could downregulate the protein Thrombospondin2 (THBS2) and thereby inhibit migration in breast cancer. (Qi et al., 2023). In pancreatic cancer the protein Menin was reported to inhibit EMT and metastasis when C/EBPβ was available. In the absence of C/EBPβ, Menin acted as an oncogene and supported cellular motility and EMT (Cheng et al., 2019). Contrary to these studies, the LIP protein isoform of C/EBPβ was shown to inhibit the transcription of miR-203, a tumor suppressive miRNA that is known to inhibit cellular metastasis. Expression of LIP in esophageal cancer cells could exacerbate EGF-induced EMT (Li et al., 2014).

Among the other transcription factors identified were RELA, which encodes the protein p65 and is a component of the master transcription factor NF-κB. NF-κB was shown to enhance both initiation and progression of EMT in many different cancer types through regulation of cellular junctions, cytoskeletal reorganization, loss of polarity and induction of expression of extracellular matrix remodelling enzymes (Oh et al., 2023). In HCC, the cytokine IL-17A was shown to activate NF-κB, which in turn activated matrix metalloproteases to induce EMT (Li et al., 2011).

## 5. Conclusions

54 different miRNAs were identified in exosomes released from HCC cells undergoing p-EMT. Among these, 13 were also identified in exosomes isolated from HCC patients and were utilized for further analysis. 11 of the 13 miRNAs were of prognostic significance, showing both unfavorable (n=7) and favorable (n=4) prognosis. The association of the seven miRNAs with worse prognosis could be validated in multiple publicly available datasets. A Bayesian probability network identified transcription factors to be the common denominator for the miRNAs, their association with prognosis and the pathways and genes they target. Overall, our data suggest that HCC cells undergoing partial EMT release exosomes containing miRNAs that are associated with both worse and favorable prognosis by regulating the transcription of genes.

## Authors contributions

OM, LNE, SBakhshi: Data analysis and interpretation, drafting, OK, Generation of Slug overexpressing cells, exosome isolation, miRNA-seq, HS: drafting, funding, supervision, ACA: supervision and drafting, SBanerjee: overall supervision, conceptualization, drafting.

## Conflict of interest and acknowledgements

The authors have no conflicts of interest to disclose. This study was supported by Anadolu University projects (1705F240, 1508F587) and Orta Doğu Teknik Üniversitesi BAP project (ÇDAP-108-2020-10197).

## Compliance with ethical standards

The authors did not perform experiments with human participants or animal models in this study.

## Abbreviations

AUROC: Area Under the Receiver Operating Characteristic Curve
DE: Differential expression
COL2A1: Collagen type II alpha 1
EMT: Epithelial to mesenchymal transition
EV: Empty vector (control)
FGG: Native fibrinogen gamma chain
FN1: Fibronectin 1
FPR: False Positive Rate
GEO: Gene Expression Omnibus
HCC: Hepatocellular carcinoma
KM: Kaplan Meier
MET: Mesenchymal to epithelial transition
MSRS: Maximally selected rank statistics
OS: Overall survival

## Notes

### Competing Interest Statement

The authors have declared no competing interest.

