## Supplemental Figure 1 for "Development of an EMT-related exosomal miRNA signature that can predict prognosis in hepatocellular carcinoma": Supplementary Figure 1.docx

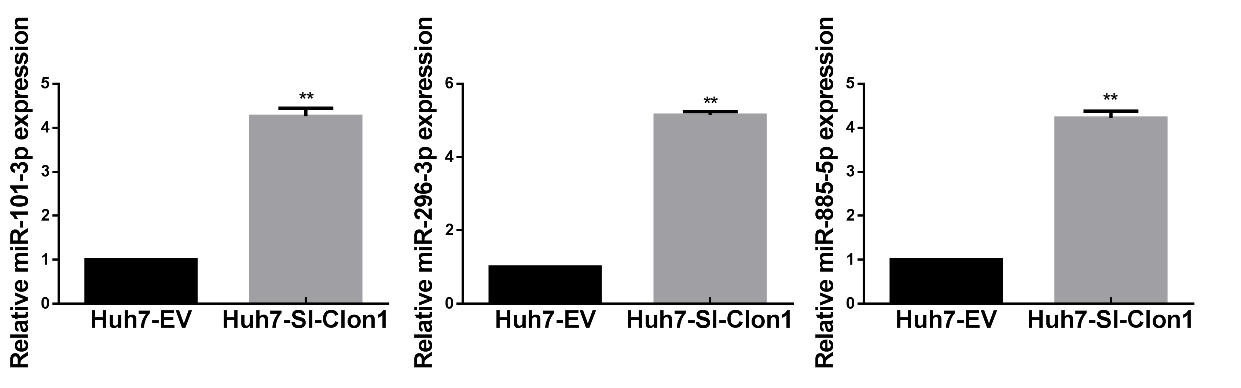

Supplementary Figure 1 Relative levels of miR-101-3p, miR-296-3p and miR-885-5p in control and (Huh7-EV) and Slug overexpressing (Huh7-SI-Clon1) HuH7 cells. The experiments were performed as three independent biological replicates. Statistical comparisons were carried out using Student t-test (**p < 0.01).
