## Supplemental Figure 1 for "Development of an EMT-related exosomal miRNA signature that can predict prognosis in hepatocellular carcinoma": Supplementary Table 4.docx

Supplementary Table 4 Fifty-six miRNAs that are upregulated in exosomes isolated from non-benign HCC tumors versus exosomes collected from benign tumors ([GSE209611](https://www.ncbi.nlm.nih.gov/geo/query/acc.cgi?acc=GSE209611))

| **miRBase Accession*** | **FDR** | **p-value** | **t-value** | **logFC** | **miRNA** |
| --- | --- | --- | --- | --- | --- |
| MIMAT0015052 | 8.43E-03 | 3.71E-03 | 3.50031 | 1.3269 | hsa-miR-3175 |
| MIMAT0028125 | 1.09E-02 | 4.88E-03 | 3.35993 | 1.18523 | hsa-miR-7114-5p |
| MIMAT0016915 | 1.10E-02 | 4.92E-03 | 3.35594 | 1.82275 | hsa-miR-4284 |
| MIMAT0019032 | 1.31E-02 | 5.92E-03 | 3.26125 | 1.41938 | hsa-miR-4497 |
| **MIMAT0000267** | **1.99E-02** | **9.18E-03** | **3.03745** | **1.82775** | **hsa-miR-210-3p** |
| MIMAT0000093 | 2.03E-02 | 9.34E-03 | 3.02846 | 1.01514 | hsa-miR-93-5p |
| MIMAT0027464 | 2.46E-02 | 1.15E-02 | 2.92121 | 1.40469 | hsa-miR-6782-5p |
| MIMAT0018929 | 2.57E-02 | 1.20E-02 | 2.89855 | 1.78713 | hsa-miR-4417 |
| MIMAT0018085 | 2.71E-02 | 1.27E-02 | 2.86977 | 1.203 | hsa-miR-3663-3p |
| MIMAT0028211 | 2.77E-02 | 1.30E-02 | 2.85811 | 2.16317 | hsa-miR-7150 |
| **MIMAT0000081** | **3.01E-02** | **1.42E-02** | **2.81287** | **1.97616** | **hsa-miR-25-3p** |
| MIMAT0027383 | 3.12E-02 | 1.48E-02 | 2.793 | 1.11149 | hsa-miR-6741-5p |
| MIMAT0027620 | 3.13E-02 | 1.48E-02 | 2.79056 | 1.90357 | hsa-miR-6769b-5p |
| **MIMAT0005891** | **3.29E-02** | **1.57E-02** | **2.76312** | **1.6821** | **hsa-miR-1303** |
| MIMAT0019743 | 3.46E-02 | 1.65E-02 | 2.7365 | 1.71666 | hsa-miR-4667-5p |
| MIMAT0000076 | 3.47E-02 | 1.65E-02 | 2.73548 | 2.24231 | hsa-miR-21-5p |
| **MIMAT0000728** | **3.51E-02** | **1.68E-02** | **2.7283** | **1.86037** | **hsa-miR-375** |
| MIMAT0027686 | 3.97E-02 | 1.90E-02 | 2.66217 | 1.60079 | hsa-miR-6893-5p |
| **MIMAT0000099** | **4.14E-02** | **1.99E-02** | **2.63888** | **1.55492** | **hsa-miR-101-3p** |
| MIMAT0027690 | 4.23E-02 | 2.05E-02 | 2.62481 | 1.3816 | hsa-miR-6895-5p |
| MIMAT0018178 | 4.87E-02 | 2.39E-02 | 2.54316 | 1.2193 | hsa-miR-3180 |
| MIMAT0003945 | 5.03E-02 | 2.48E-02 | 2.52538 | 1.56403 | hsa-miR-765 |
| MIMAT0019878 | 5.06E-02 | 2.50E-02 | 2.52086 | 1.26648 | hsa-miR-4745-5p |
| **MIMAT0000272** | **5.20E-02** | **2.57E-02** | **2.50589** | **1.66195** | **hsa-miR-215-5p** |
| MIMAT0019838 | 5.47E-02 | 2.71E-02 | 2.4774 | 1.43808 | hsa-miR-4723-5p |
| MIMAT0022943 | 5.49E-02 | 2.73E-02 | 2.47447 | 1.76446 | hsa-miR-1233-5p |
| MIMAT0027572 | 5.68E-02 | 2.83E-02 | 2.45503 | 1.83422 | hsa-miR-6780b-5p |
| MIMAT0025476 | 5.68E-02 | 2.83E-02 | 2.45481 | 1.29342 | hsa-miR-6510-5p |
| MIMAT0022709 | 5.70E-02 | 2.84E-02 | 2.45336 | 1.78681 | hsa-miR-652-5p |
| MIMAT0022942 | 6.02E-02 | 3.03E-02 | 2.42034 | 1.43677 | hsa-miR-1229-5p |
| **MIMAT0004692** | **6.03E-02** | **3.03E-02** | **2.41933** | **1.42881** | **hsa-miR-340-5p** |
| MIMAT0022721 | 6.19E-02 | 3.12E-02 | 2.40435 | 1.89409 | hsa-miR-1247-3p |
| MIMAT0018981 | 6.38E-02 | 3.22E-02 | 2.38779 | 2.09265 | hsa-miR-4459 |
| MIMAT0027658 | 6.48E-02 | 3.28E-02 | 2.37764 | 1.42536 | hsa-miR-6879-5p |
| MIMAT0018068 | 6.48E-02 | 3.28E-02 | 2.3776 | 1.30059 | hsa-miR-3648 |
| **MIMAT0004947** | **6.59E-02** | **3.34E-02** | **2.36768** | **1.41052** | **hsa-miR-885-5p** |
| MIMAT0015079 | 6.63E-02 | 3.37E-02 | 2.36404 | 1.23211 | hsa-miR-3195 |
| MIMAT0019868 | 6.82E-02 | 3.47E-02 | 2.34773 | 1.57924 | hsa-miR-4739 |
| MIMAT0019885 | 6.83E-02 | 3.48E-02 | 2.34644 | 1.18189 | hsa-miR-4749-5p |
| **MIMAT0000074** | **6.85E-02** | **3.49E-02** | **2.3445** | **1.19471** | **hsa-miR-19b-3p** |
| MIMAT0019022 | 6.95E-02 | 3.54E-02 | 2.33662 | 1.40466 | hsa-miR-4488 |
| MIMAT0009448 | 7.37E-02 | 3.77E-02 | 2.30359 | 1.65918 | hsa-miR-1973 |
| MIMAT0020924 | 7.44E-02 | 3.81E-02 | 2.29755 | 1.96293 | hsa-miR-642a-3p |
| MIMAT0016878 | 7.51E-02 | 3.86E-02 | 2.29161 | 1.1127 | hsa-miR-4257 |
| MIMAT0019964 | 7.64E-02 | 3.93E-02 | 2.28135 | 1.50616 | hsa-miR-4792 |
| MIMAT0000073 | 7.86E-02 | 4.06E-02 | 2.26443 | 1.263 | hsa-miR-19a-3p |
| MIMAT0002877 | 8.15E-02 | 4.23E-02 | 2.24271 | 1.62844 | hsa-miR-513a-5p |
| MIMAT0002816 | 8.17E-02 | 4.25E-02 | 2.24023 | 1.57289 | hsa-miR-494-3p |
| MIMAT0000101 | 8.26E-02 | 4.30E-02 | 2.23355 | 1.02309 | hsa-miR-103a-3p |
| MIMAT0019043 | 8.49E-02 | 4.42E-02 | 2.21818 | 1.49729 | hsa-miR-2392 |
| MIMAT0027363 | 8.63E-02 | 4.51E-02 | 2.20733 | 1.43436 | hsa-miR-6731-5p |
| **MIMAT0000275** | **8.97E-02** | **4.70E-02** | **2.1855** | **1.15903** | **hsa-miR-218-5p** |
| MIMAT0019852 | 9.31E-02 | 4.89E-02 | 2.16389 | 1.45352 | hsa-miR-4730 |
| **MIMAT0005920** | **9.36E-02** | **4.93E-02** | **2.16024** | **1.40638** | **hsa-miR-1266-5p** |
| **MIMAT0004679** | **9.41E-02** | **4.96E-02** | **2.1564** | **1.19495** | **hsa-miR-296-3p** |

* miRNAs common with Supplementary Table 3 are shown in bold.
