## Supplemental Figure 1 for "Development of an EMT-related exosomal miRNA signature that can predict prognosis in hepatocellular carcinoma": Supplementary Table 5.docx

Supplementary Table 5: List of 81 genes identified by enrichment analysis

| **Gene Symbol** | **p-value** | **FDR** | **Odds ratio** | **Number of interactions** |
| --- | --- | --- | --- | --- |
| **BCL2L11** | **1.99E-08** | **0.0000387** | **0.0763** | **8** |
| DEAF1 | 1.31E-08 | 0.0000387 | 0.00875 | 4 |
| GIT2 | 4.68E-08 | 0.0000609 | 0.0113 | 4 |
| MACF1 | 0.000000263 | 0.000205 | 0.0163 | 4 |
| CUL5 | 0.000000865 | 0.000345 | 0.0213 | 4 |
| DENND6A | 0.000000893 | 0.000345 | 0.041 | 5 |
| **DICER1** | **0.000000691** | **0.000345** | **0.0625** | **6** |
| KAT2B | 0.000000865 | 0.000345 | 0.0213 | 4 |
| **MLEC** | **0.0000018** | **0.000582** | **0.0734** | **6** |
| WDR1 | 0.00000215 | 0.000599 | 0.0263 | 4 |
| PTP4A2 | 0.00000247 | 0.000641 | 0.05 | 5 |
| ELL2 | 0.00000364 | 0.000821 | 0.054 | 5 |
| MBNL2 | 0.00000379 | 0.000821 | 0.03 | 4 |
| **SLC7A11** | 0.00000409 | 0.000839 | 0.0842 | 6 |
| ITPR1 | 0.0000053 | 0.001 | 0.0325 | 4 |
| BTBD7 | 0.00000722 | 0.00128 | 0.035 | 4 |
| ABHD17C | 0.00000961 | 0.00156 | 0.0375 | 4 |
| PAPD7 | 0.00000924 | 0.00156 | 0.065 | 5 |
| MITF | 0.000011 | 0.00172 | 0.0388 | 4 |
| ZIC5 | 0.0000292 | 0.00345 | 0.082 | 5 |
| FRS2 | 0.000031 | 0.00355 | 0.083 | 5 |
| SATB1 | 0.000038 | 0.00389 | 0.0525 | 4 |
| AGPAT5 | 0.0000417 | 0.0039 | 0.0538 | 4 |
| MET | 0.0000417 | 0.0039 | 0.0538 | 4 |
| MYCN | 0.0000417 | 0.0039 | 0.0538 | 4 |
| NUFIP2 | 0.000045 | 0.00399 | 0.206 | 4 |
| DEPDC1 | 0.0000649 | 0.00473 | 0.06 | 4 |
| PRRC2A | 0.0000764 | 0.00497 | 0.0625 | 4 |
| CAB39 | 0.0000826 | 0.00511 | 0.0638 | 4 |
| FNDC3A | 0.0000826 | 0.00511 | 0.0638 | 4 |
| KCTD10 | 0.0000826 | 0.00511 | 0.0638 | 4 |
| ITGB1 | 0.0000963 | 0.00569 | 0.0663 | 4 |
| ACTB | 0.000107 | 0.00578 | 0.107 | 5 |
| FBXW7 | 0.000138 | 0.00578 | 0.0725 | 4 |
| RNF44 | 0.000138 | 0.00578 | 0.0725 | 4 |
| SUZ12 | 0.000128 | 0.00578 | 0.0713 | 4 |
| UBN2 | 0.000127 | 0.00578 | 0.153 | 6 |
| DDX6 | 0.000157 | 0.00638 | 0.116 | 5 |
| SERTAD2 | 0.000157 | 0.00638 | 0.075 | 4 |
| RAB8B | 0.000168 | 0.00654 | 0.0763 | 4 |
| ZFYVE26 | 0.000168 | 0.00654 | 0.0763 | 4 |
| DCBLD2 | 0.000203 | 0.00742 | 0.08 | 4 |
| GPR157 | 0.000203 | 0.00742 | 0.08 | 4 |
| HOXA9 | 0.000203 | 0.00742 | 0.08 | 4 |
| ATXN1 | 0.000334 | 0.00891 | 0.136 | 5 |
| PPP2R5E | 0.000321 | 0.00891 | 0.09 | 4 |
| CAPZB | 0.000357 | 0.00909 | 0.0925 | 4 |
| CFL2 | 0.000357 | 0.00909 | 0.0925 | 4 |
| SLC38A2 | 0.000357 | 0.00909 | 0.0925 | 4 |
| JARID2 | 0.000395 | 0.00988 | 0.095 | 4 |
| NRAS | 0.000395 | 0.00988 | 0.095 | 4 |
| RAP2C | 0.000395 | 0.00988 | 0.095 | 4 |
| ZNF772 | 0.000581 | 0.013 | 0.105 | 4 |
| RHOA | 0.000789 | 0.0151 | 0.114 | 4 |
| DYNC1LI2 | 0.000893 | 0.0163 | 0.118 | 4 |
| KLF10 | 0.00101 | 0.0175 | 0.121 | 4 |
| TGOLN2 | 0.00113 | 0.0191 | 0.125 | 4 |
| ARID1A | 0.00122 | 0.0192 | 0.128 | 4 |
| SZRD1 | 0.00117 | 0.0192 | 0.126 | 4 |
| UBE2D3 | 0.00151 | 0.021 | 0.135 | 4 |
| TGFBR2 | 0.00162 | 0.0223 | 0.138 | 4 |
| WNK1 | 0.00162 | 0.0223 | 0.138 | 4 |
| OTUD7B | 0.00173 | 0.0232 | 0.14 | 4 |
| AGO1 | 0.00185 | 0.0241 | 0.143 | 4 |
| FZD6 | 0.00191 | 0.0247 | 0.144 | 4 |
| RORA | 0.00203 | 0.0253 | 0.146 | 4 |
| DDX3X | 0.0021 | 0.0259 | 0.148 | 4 |
| MBNL1 | 0.0023 | 0.0278 | 0.151 | 4 |
| SON | 0.0023 | 0.0278 | 0.151 | 4 |
| MYC | 0.00238 | 0.0282 | 0.153 | 4 |
| MAP3K9 | 0.00245 | 0.0288 | 0.154 | 4 |
| TP53 | 0.003 | 0.0328 | 0.163 | 4 |
| CCDC80 | 0.00345 | 0.0357 | 0.169 | 4 |
| DSN1 | 0.00426 | 0.0357 | 0.179 | 4 |
| IGF1R | 0.00488 | 0.0357 | 0.245 | 5 |
| LZIC | 0.00345 | 0.0357 | 0.169 | 4 |
| XIAP | 0.0043 | 0.0357 | 0.238 | 5 |
| ZBTB18 | 0.00556 | 0.0378 | 0.193 | 4 |
| PKM | 0.00638 | 0.0417 | 0.2 | 4 |
| SOX4 | 0.0076 | 0.0464 | 0.21 | 4 |
| PPP1R15B | 0.00792 | 0.0472 | 0.213 | 4 |
